# Information alignment between interacting brains

**DOI:** 10.1101/2025.01.07.631802

**Authors:** Denise Moerel, Tijl Grootswagers, Genevieve L. Quek, Sophie Smit, Manuel Varlet

## Abstract

Social interactions shape our perception of the world, influencing how we interpret incoming information. Alignment between interacting individuals’ sensory and cognitive processes is key to successful cooperation and communication, but the neural processes underlying this alignment remain unknown. Here, we leveraged Representational Similarity Analysis (RSA) on electroencephalography (EEG) hyperscanning techniques to investigate information alignment in 24 pairs of participants who performed a categorisation task together based on agreed upon rules. Significant interbrain information alignment emerged within 45 ms of stimulus presentation and persisted for hundreds of milliseconds. Early alignment (45–180 ms) occurred in both real and randomly matched pseudo-pairs, reflecting shared sensory responses. Importantly, alignment after 200 ms strengthened with practice and was unique to real pairs, driven by shared representations associated with, and extending beyond, the categorisation rules they formed. Together, these findings highlight distinct processes underpinning interbrain information alignment during social interactions, that can be effectively captured and disentangled with Interbrain RSA.

## 1. INTRODUCTION

The groups we belong to can significantly shape our perception and interpretation of the world around us. Whether these groups are based on culture, profession, social circles, or any other common interest, they provide a shared context, set of values, and ways of interpreting incoming information, which can facilitate communication and joint activities across our society (Cialdini & Goldstein, 2004; Moore et al., 2014; Tomasello et al., 2005). Even though visual perception is largely automatic and seemingly objective (Koenig-Robert et al., 2024; Nastase et al., 2019; Robinson et al., 2023), the interpretation of visual information can be strongly influenced by social interactions, modulating how similarly people represent and respond to their environment depending on whether they belong to same or different groups (Cialdini & Goldstein, 2004; Coey et al., 2012; Sebanz et al., 2006). Hyperscanning, a technique that allows for the simultaneous measurement of brain activity in multiple people while they interact with each other, has become a valuable tool for studying real-time social interactions and the dynamics of interacting brains (Czeszumski et al., 2020; Dumas et al., 2010; Zamm et al., 2024). While previous hyperscanning research has focused on interbrain synchrony measures, recent evidence suggests that these measures are content agnostic and do not capture information alignment across people, and that multivariate representational analyses are a powerful alternative (Moerel et al., 2025; Varlet & Grootswagers, 2024; Zada et al., 2024). Here we leverage and advance EEG hyperscanning methods to uncover how representations in interacting individuals’ brains evolve in real-time and uniquely converge as a function of commonly agreed upon rules and interpretation of their visual environment.

Previous hyperscanning research has revealed enhanced synchrony between the neural oscillations of individuals as they engage in joint activities, which has been argued to support communication, coordination and collaboration (for an overview, see Czeszumski et al., 2020; Kelsen et al., 2022; Mu et al., 2018; Shemyakina & Nagornova, 2021). Higher neural alignment between people, as indexed by a range of measures of synchrony (e.g., Coherence, Phase Locking Value, weighted Phase Lag Index), has been shown during genuine social interactions – participants interacting together in real-time vs. participants being paired a posteriori (pseudo pairs; Pick et al., 2024; Reindl et al., 2022) – and with higher levels of familiarity (e.g., mother-infant interaction; Endevelt-Shapira et al., 2021; Kinreich et al., 2017) and collaborations (e.g., collaborative vs. competitive scenarios; Babiloni et al., 2007; Fallani et al., 2010; Sinha et al., 2016). However, several recent studies have questioned the nature of the processes captured by interbrain synchrony measures (Burgess, 2013; Hamilton, 2021; Holroyd, 2022; Novembre & Iannetti, 2021), and revealed critical methodological limitations (Burgess, 2013; Varlet & Grootswagers, 2024; Zimmermann et al., 2024).

One study found evidence for coincidental synchrony between pseudo pairs using Phase-Locking Value and to a lesser degree Coherence and Partial Directed Coherence measures (Burgess, 2013). This coincidental synchrony has been attributed to common EEG rhythmicity across participants, arising from systematic differences between experimental conditions. Another study provided evidence that methodological decisions, such as the use of short epochs, can inflate phase-based estimates of interbrain synchrony (Zimmermann et al., 2024). Finally, recent work highlighted that phase- as well as amplitude-based measures of interbrain synchrony were not sensitive to the alignment of *information* across people, as they were minimally affected by whether participants saw the same or different objects (Varlet & Grootswagers, 2024). The authors introduced Interbrain Representational Similarity Analysis (RSA), an adaption of RSA (Kriegeskorte et al., 2008), to more effectively capture and compare the information content represented in each brain. Unlike synchrony measures, Interbrain RSA is information-based, allowing researchers to identify what information is shared between individuals at specific moments in time. It also provides a single, experiment-level estimate of information alignment between brains, making it less susceptible to transient fluctuations driven by shared environmental factors, such as changes in lighting or acoustic conditions. Finally, by abstracting away from the raw data and focusing on the representational structure contained in the data, Interbrain RSA enables comparisons across different brain regions, time scales, neuroimaging modalities and tasks.

Interbrain RSA holds promise to capture and understand how visual representations evolve and converge across individuals through social interactions. However, its capacity to uncover information alignment beyond that simply driven by robust differences in evoked sensory responses, as modulated by same vs. different visual objects in Varlet and Grootswagers (2024), remains to be demonstrated. Indeed, understanding how the same visual stimuli are (dis)similarly represented by individuals depending on the groups they belong to critically requires not only being able to track information alignment that is reliably and externally evoked, but also alignment that is socially learnt and internally induced (Holroyd, 2022). The latter type of information alignment is expected to reflect convergence in higher-level representations, extending beyond initial visual responses occurring as early as ∼50 ms after stimulus onset (Carlson et al., 2013; Cichy et al., 2014; Contini et al., 2017), and engaging more distributed brain networks involved in attention and decision-making. Consistent with this notion, previous studies have demonstrated task- and decision-relevant modulations of visual information processing from 140-220 ms onwards (Bode et al., 2012; Grootswagers et al., 2021; Moerel et al., 2022, 2024; Quek & de Heering, 2024) and within frontal-parietal networks (Jackson et al., 2016; Jackson & Woolgar, 2018; Moerel et al., 2024; Woolgar et al., 2015). Regions across the frontal cortex, including the anterior cingulate and medial prefrontal cortices, have also been implicated in hyperscanning research during real-time social interactions (Babiloni et al., 2007).

In this study, we used Interbrain RSA with EEG hyperscanning to investigate the emergence and dynamics of information alignment between interacting individuals’ brains while presented with various visual stimuli. We aimed to distinguish information alignment that was purely evoked from participants seeing the same thing at the same time, from alignment that was driven by each pair’s pre-formed rules and interpretations. We recorded EEG data from 24 pairs of participants sitting back-to-back while performing a 4-way categorisation task on visual stimuli. Importantly, the two participants within each pair had previously created the 4 groups together by agreeing on arbitrary categorisation rules (see Figure 1). We used Interbrain RSA to determine the time-course of neural information alignment between individuals within a pair during the categorisation task. To determine whether and when the neural information alignment was purely evoked by shared visual information (seeing the same stimulus), we compared Interbrain RSA between real participant pairs and randomly matched-up pseudo pairs who saw the same stimuli in the same order. Finally, we used randomly matched-up pseudo pairs who independently created the same 4 groups of stimuli to determine the extent to which the alignment was unique to the pair or driven by the pair’s pre-formed rules. We expected alignment, beyond that driven by shared visual input, to be reflected by higher Interbrain RSA in real pairs, which would occur during later information processing stages, be more widely distributed across the brains, and strengthen over time with practice and integration of pre-agreed categorisation rules. Moreover, we examined whether this neural information alignment persisted beyond the categorisation task, which we tested by recording EEG data during passive viewing of the same stimuli in a rapid visual stream before and after participants agreed on the categorisation rules and performed the categorisation task.

**Figure 1.**
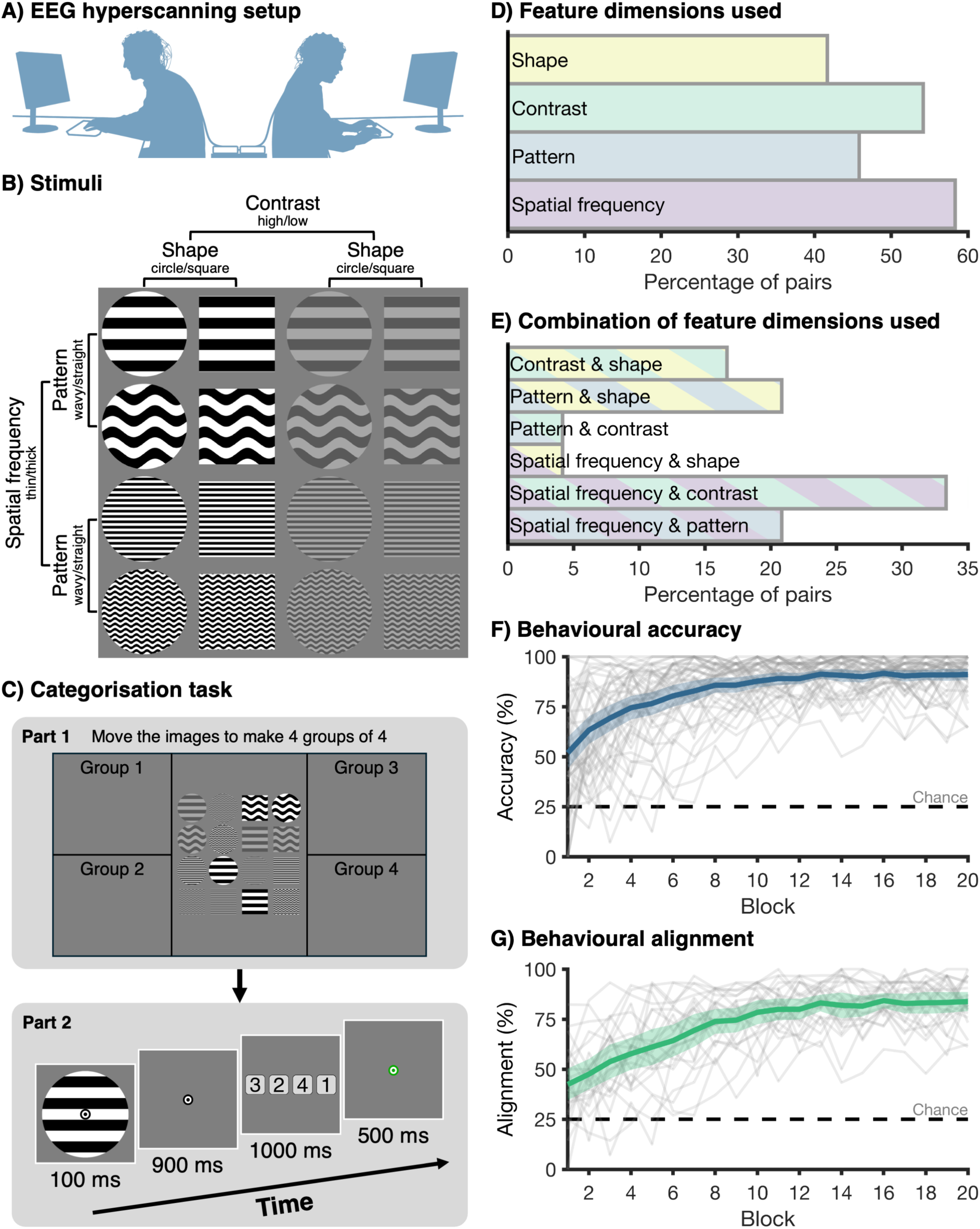
Stimuli, experiment procedure, and behavioural results. **A)** Participants were seated back-to-back, facing separate but identical displays. Each participant had a keyboard to provide responses. **B)** There were 16 unique greyscale stimuli, varying on 4 dimensions: spatial frequency (thick vs. thin lines), pattern (wavy vs. straight lines), contrast (high vs. low) and general shape (circle vs. square). **C)** The two-part categorisation task. In Part 1 (top panel), participants worked together to sort the stimuli into 4 groups of 4 stimuli. To do this, they had to select two dimensions to group the stimuli, while ignoring the two other dimensions. In Part 2 (bottom panel), participants categorised stimuli into the 4 groups they just created: Participants saw a stimulus for 100 ms, followed by a blank screen for 900 ms. Then, a response-button mapping screen was shown for 1000 ms, and participants could make a categorisation response based on the 4 groups that were made in Part 1. This was followed by a feedback screen for 500 ms, providing feedback about whether the response of the two participants matched the previously agreed upon sorting rules. The feedback pertained to the pair, not the individual. During the feedback, the fixation bullseye turned green if both individuals in the pair were correct, red if neither individual was correct, and orange if only one individual was correct. In the latter case, the feedback was not specific about which individual in the pair was correct. **D)** The features that were used by the pairs to make the 4 groups. The x-axis shows the percentage of pairs that chose each feature. **E)** The combinations of features that were used to make the 4 groups. Each pair of participants chose two features. The x-axis shows the percentage of pairs that chose each combination of features. **F)** Behavioural accuracy on the categorisation task over experiment blocks. The thick blue line shows the average across the 48 individuals. The shaded blue area shows the 95% confidence interval. The thin grey lines show the accuracy of individual participants. Chance is at 25% in this 4-way categorisation task (dotted line). **G)** Response alignment between the two participants in the pair, regardless of whether the response was correct. The green line shows the average alignment across the 24 pairs. The shaded green area shows the 95% confidence interval, and the dotted line indicates chance level (25%).

## 2. RESULTS

### 2.1. Commonly agreed rules and behavioural alignment

We recorded the brain activity of 24 pairs of participants using the 64 channel BioSemi active electrode system (BioSemi, Amsterdam, The Netherlands). The two participants in the pair were seated back-to-back in the same dimly lit room, viewing separate computer screens that showed identical visual input throughout the task (Figure 1A). Participants engaged in a joint categorisation task, which consisted of two parts. In the first part, the two participants in the pair agreed upon sorting rules for visual stimuli, and in the second part, participants used these shared rules to categorise the images. There were 16 images that varied across 4 dimensions, with 2 levels per dimension (Figure 1B): spatial frequency (thick vs. thin lines), pattern (wavy vs. straight lines), contrast (high vs. low) and general shape (circle vs. square). During the first part of the task, the two participants in the pair worked together to sort the images into 4 groups of 4, effectively selecting two of the stimulus dimensions while ignoring the other two dimensions (Figure 1C, top panel). Participants were allowed to talk during this stage. They saw identical screens and shared a cursor, which means they could both drag and drop the stimuli and observe the movements of the other participant. Both participants had to agree on the arbitrary rules before continuing the task. To determine whether the different pairs came up with different strategies, we analysed which rules were used by each pair (Figure 1D). All dimensions were used across the 24 pairs: spatial frequency was chosen most often (58.33% of pairs), followed by contrast (54.17% of pairs), pattern (45.83% of pairs), and shape (41.67% of pairs). We also assessed which *combination* of two dimensions was used by the pairs (Figure 1E). Again, all combinations of dimensions were used, with the largest group choosing spatial frequency and contrast (33.33% of pairs), followed by spatial frequency and pattern as well as pattern and shape (20.83% of pairs), contrast and shape (16.67% of pairs), and by spatial frequency and shape as well as pattern and contrast (4.16% of pairs). These results show that all dimensions were used across pairs as classification rules, with a reasonable spread across all options. This means that the stimulus sorting rules were indeed arbitrary, in that different pairs of individuals derived different sets of rules during the communication phase.

During the second part of the categorisation task, the participants categorised the stimuli based on the rules they had just agreed upon by allocating each stimulus to one of the four newly-defined groups (Figure 1C, bottom panel). Participants would see a stimulus for 100 ms, followed by a blank screen for 900 ms, during which participants could make a decision about which category the stimulus belonged to. After the blank screen, four button responses were shown for 1000 ms, which both participants had to select using four different keys on their keyboard to provide a categorisation response. The mapping between the four buttons and four categories was randomly changed every trial. This means participants could not know which key to press before the response mapping was presented on the screen. This delayed response screen with unpredictable button mappings ensured that the 1000 ms of EEG data after stimulus onset – our window of interest – were free from any motor preparation and contamination. Participants were asked not to talk during this part of the task, to avoid contamination of the EEG signal, but they were allowed to talk during the self-paced breaks.

Breaks occurred after every 32 trials, with less than 3 minutes between two consecutive breaks, and there were a total of 19 breaks throughout the task.

We calculated the behavioural accuracy at the level of the individual (Figure 1F), by determining the percentage of responses that matched with the previously agreed upon sorting rules. In addition, we calculated the alignment in behavioural responses between the two individuals in the pair, by determining whether the two individuals in the pair provided the same response, regardless of whether it was correct or incorrect (Figure 1G). Visual inspection reveals an upward trend for both behavioural measures, with higher accuracy as well as higher response alignment later in the experiment.

### 2.2. Evidence for information alignment beyond shared visual input

We used Interbrain RSA to assess the degree to which neural representations were aligned between the two individuals in the pair during the categorisation task. For each participant separately, we created a Representational Dissimilarity Matrix (RDM) for each time-point in the epoch, reflecting the dissimilarity in the pattern of activation across all 64 channels for each combination of two unique stimuli. To determine the time-course of information alignment between the two individuals in the pair, accounting for information alignment at different time-points between individuals, we calculated the correlation between the dissimilarity matrices of the two individuals for each combination of time-points in the epoch. Figure 2 (left panel) shows the information alignment between the participants in a pair, averaged across pairs. To determine whether the observed information alignment was due to simply viewing the same visual input, we obtained the same measure for randomly matched pseudo pairs, based on 10,000 random pair assignments. Importantly, we matched the counterbalancing of the trial orders for the pseudo pairs, with 4 unique trial orders among the 24 pairs. This ensured that the individuals in the pseudo pairs saw the same stimuli, in the same order, as the real pairs. Figure 2 (right panel) shows the information alignment for the pseudo pairs. Visual inspection shows a remarkably similar pattern of strong information alignment early in time, from approximately 50 to 165 ms after stimulus onset. However, the information alignment appears more sustained over time for the real pairs compared to the randomly matched pseudo pairs.

**Figure 2.**
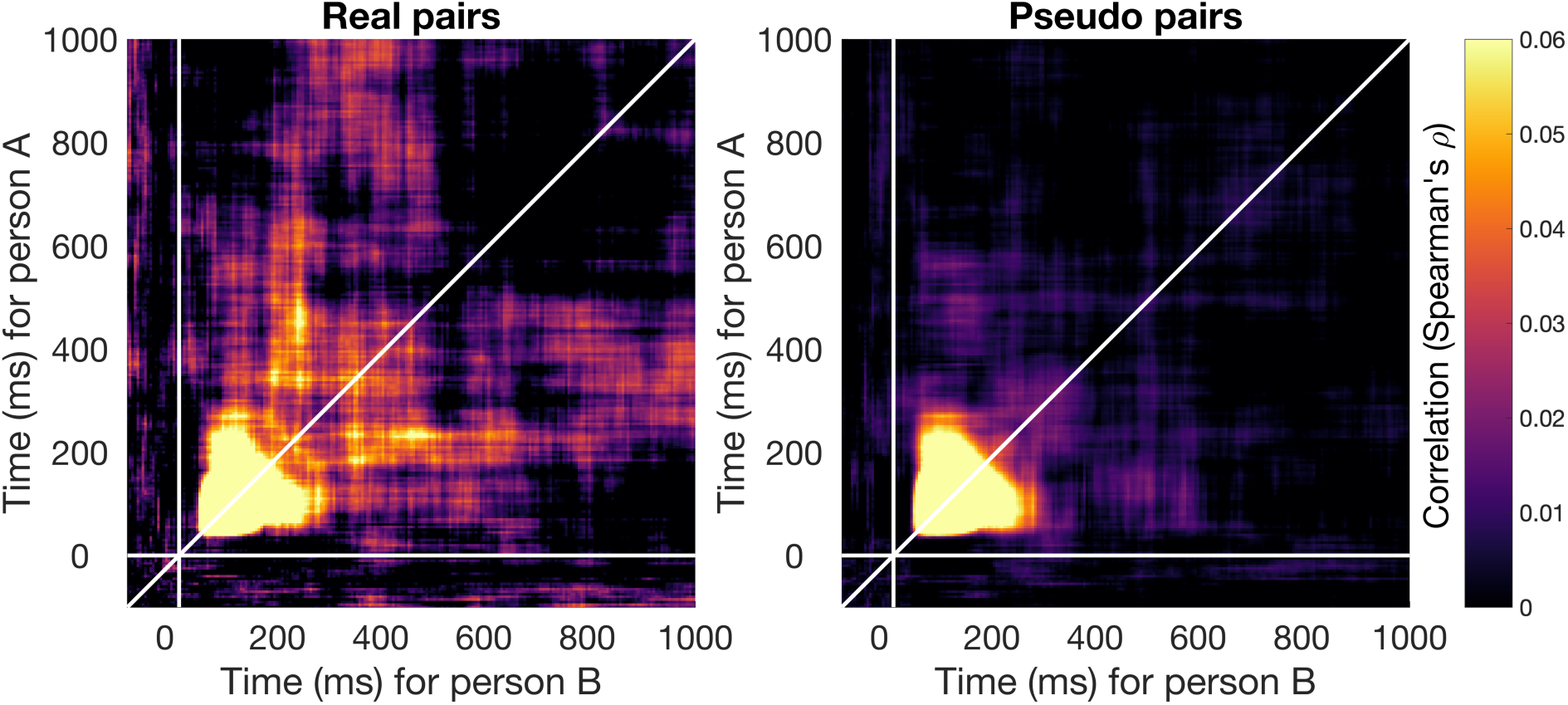
Information alignment between the two individuals in each pair. The left panel shows the mean (Spearman) correlation between the neural representation of the two individuals in each real pair for all combinations of time-points. The brightness of the colour reflects the strength of the correlation, with brighter colours reflecting higher correlations, and therefore stronger information alignment. The y-axis shows the time for person A in the pair, and the x-axis the time for person B in the pair. The right panel shows the information alignment for the pseudo pairs, who did not perform the task together. Here, we repeated the same analysis for randomly matched pseudo pairs instead of real pairs and with results showing the average across 10,000 iterations.

To reduce dimensionality and obtain a single time-course of information alignment, we averaged over one time dimension of the symmetrical time-by-time matrix (Figure 2), enabling us to capture information alignment across all possible time lags between the two participants (Figure 3A). The time-course shows the alignment for a given time-point of person A with all other time-points for person B in the pair and vice versa. Cluster-based permutation testing revealed significant alignment in the real pairs from 45 to 285 ms and 315 to 425 ms after stimulus onset. This means that during this time, the information alignment between individuals in a pair was stronger than would be expected by chance. There was no significant difference between the real pairs and randomly matched pseudo pairs, who saw the same stimuli in the same order, from 45 to 180 ms (no significant clusters and all p values > .05). This suggests that early in time, the information alignment between participants was driven by shared visual information. However, there was a significant difference between the real pairs and randomly matched pseudo pairs from 185 to 240 ms, 325 to 360 ms, and 370 to 425 ms after stimulus onset, with stronger information alignment observed for the real pairs compared to pseudo pairs. This suggests that later in time, there was information alignment between individuals in the real pairs that was not purely visually evoked but uniquely emerged from commonly agreed upon rules and interpretations.

**Figure 3.**
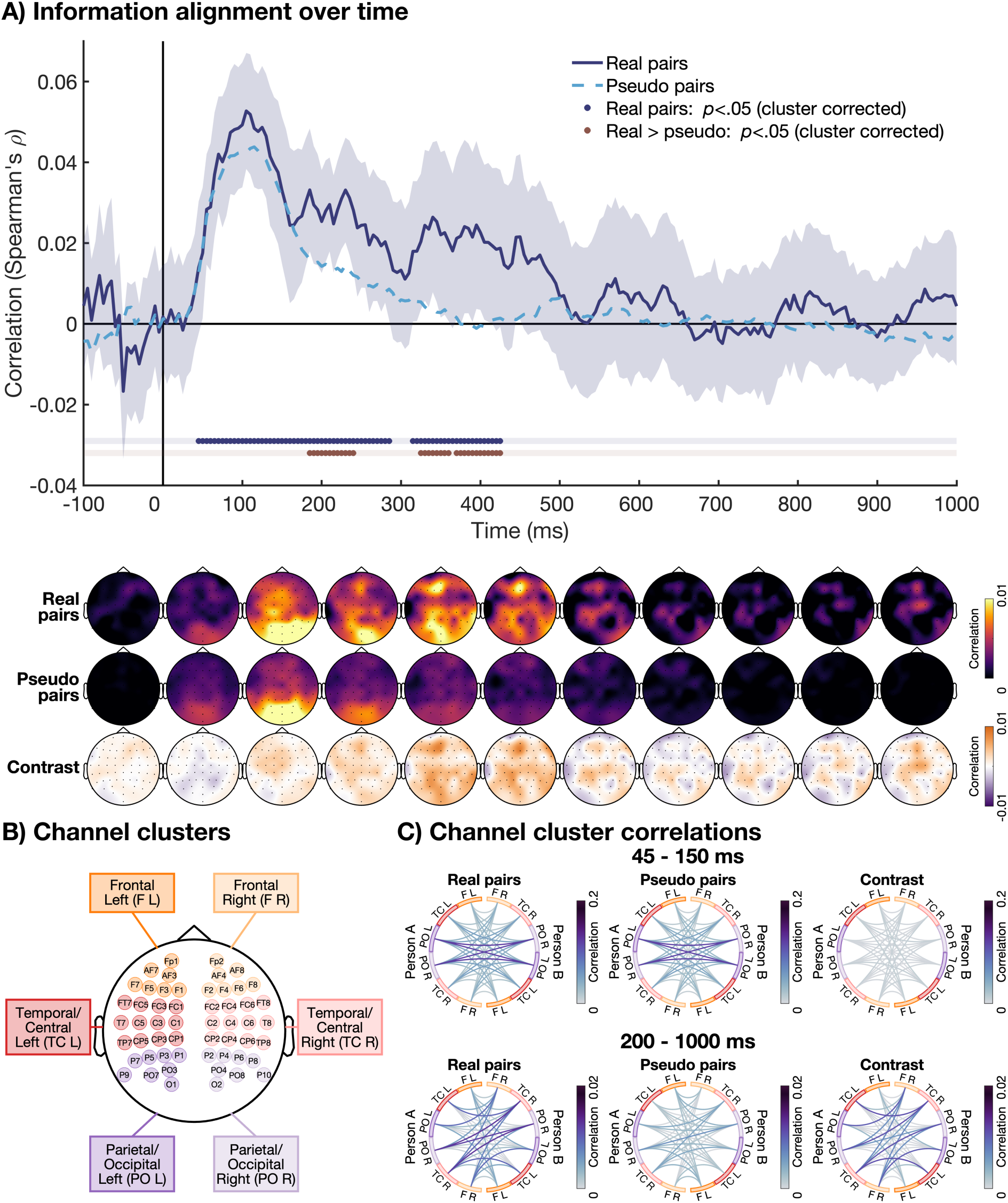
The time-course of information alignment. **A)** The time-course of information alignment is shown in dark blue. This measure was obtained by collapsing the temporal generalisation matrix of dissimilarity matrix correlations between the two individuals in each pair (Figure 2) along one of the time dimensions. Cluster corrected p values < .05 for this correlation are shown in dark blue below. The shaded area shows the 95% confidence interval. The time-course of dissimilarity matrix correlations for the randomly matched pseudo pairs is shown as a dashed light blue line. This shows the information alignment that is driven by sensory evoked signals. Cluster corrected p values < .05 for the difference between the real and pseudo pairs are shown in red below. This shows the information alignment that was not driven by sensory evoked signals. The topographies for the information alignment are shown below the plot. They show the average correlation of one channel with all channels or the other participant, averaged across 100 ms time bins. The top row shows the topographies for the real pairs, the middle row for the pseudo pairs, and the bottom row for the contrast (real – pseudo pairs). **B)** The channels included in the 6 clusters: Frontal Left (F L), Frontal Right (F R), Temporal/Central Left (TC L), Temporal/Central Right (TC R), Parietal/Occipital Left (PO L), and Parietal/Occipital Right (PO R). **C)** The dissimilarity matrix correlations between person A (left) and person B (right) in the pair, for all combinations of 6 channel clusters. Correlations are averaged across an early (45 – 150 ms) and late (200 – 1000 ms) time-window. The colour of the lines reflects the strength of the correlation, with darker colour reflecting stronger information alignment. The left plots show the correlation within real pairs, the middle plots show the correlation for the pseudo pairs, and the right plots show the difference between real and pseudo pairs.

To determine which channels were driving the information alignment between individuals in a pair, we calculated for each participant the correlation at each channel with all channels from the other participant. The dissimilarity matrices for each channel, used to obtain the time-by-time map, were computed using 4 or 5 neighbouring channels. The single time-course correlation obtained from these dissimilarity matrices was then averaged into 100 ms intervals to obtain scalp topographies. These topographies indicated for each channel and time bin of participant A, how strong the alignment was with any of the other channels at any time bin for participant B and vice versa (Figure 4A). They showed that before 100 ms, the alignment was mostly driven by posterior channels, moving to more frontal channels between 100 and 200 ms. Importantly, the alignment in real and pseudo pairs looked very similar during this time. Around 300 to 400 ms, visual inspection appeared to show stronger alignment for real compared to pseudo pairs. This later alignment was driven by channels that were distributed across the scalp.

**Figure 4.**
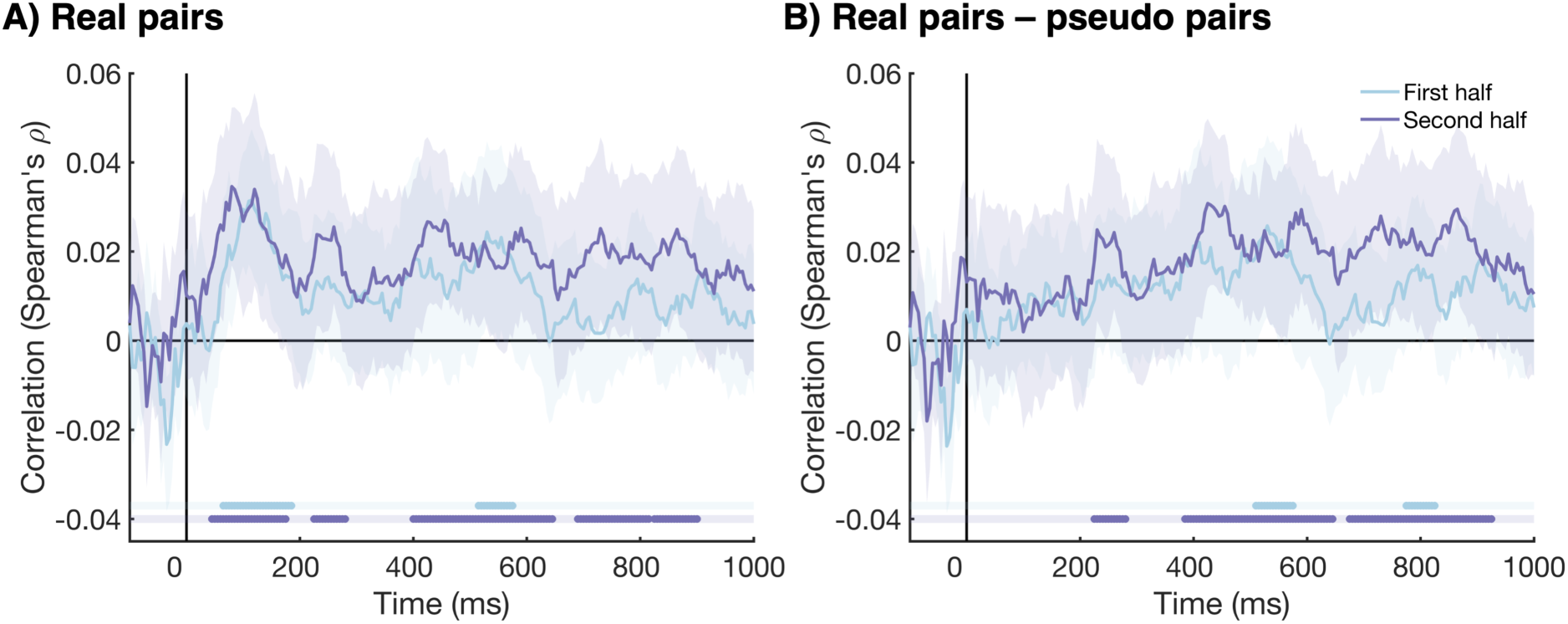
The time-course of information alignment, split by first and second experiment half. **A)** The time-course of the information alignment between the two participants in each pair for the first experiment half (light blue), and the second experiment half (purple). The shaded areas around the plot lines show the 95% confidence intervals. Cluster corrected *p* values < .05 are shown below the plot in the same colours. **B)** The difference in information alignment between the real pairs and pseudo pairs (real - pseudo). This shows the alignment that is not driven by visual information. All plotting conventions are the same as Figure 4A.

To further explore the spatial dynamics of the information alignment, and identify the most aligned regions across brains, we repeated the same analysis used for the topographies, but this time keeping the combinations of channels for both brains and grouping them into 6 pre-defined clusters to reduce dimensionalities (Figure 4B). We collapsed the time-course into early time-points (45 to 150 ms) and late time-points (200 to 1000 ms). The resulting channel cluster correlations (Figure 4C) indicated for each cluster of channels and time bin of participant A, how strong the alignment was with individual clusters of channels for participant B in the same time bin. The results show that for the early time window, there was information alignment for all clusters of channels, with the strongest alignment between left and right parietal/occipital channels (Figure 4C). Visual inspection reveals a similar pattern for the real and pseudo pairs, adding further evidence that early alignment is driven by visual information. For the late time window (200 – 1000 ms), there was stronger information alignment for real compared to pseudo pairs in channel clusters distributed across the scalp. The strongest alignment was observed between the left and right temporal/central clusters. These findings are in line with a distributed network of regions driving the later neural information alignment.

### 2.3. Late information alignment strengthens over the experiment

The behavioural categorisation data showed that participants became more accurate over the course of the experiment, with more similar responses given by the two individuals in the pairs in later blocks. We assessed whether this finding was mirrored in the information alignment between participants, by repeating the same Interbrain RSA analysis as described above separately for the first and second half of the experiment. Figure 4A shows the information alignment between participants in a pair for the two experiment halves. We found evidence for neural information alignment during the early time window for both experiment halves; between 65 and 185 ms after stimulus onset for the first experiment half and between 45 and 175 ms second experiment half. However, the information alignment in the later time window was more sustained for the second experiment half. For the first experiment half, we only found evidence for information alignment between 515 and 575 ms after stimulus onset, whereas for the second experiment half we observed information alignment between 225 and 280 ms, and for much of the time-window between 400 and 900 ms. We calculated the difference between real and pseudo pairs to assess the neural information alignment that was not driven by shared visual information (Figure 4B). The early information alignment disappeared, suggesting this alignment was simply evoked by visual information. The information alignment in the later time window was still present after subtracting the information alignment of the pseudo pairs, suggesting that the later information alignment emerged from pair’s pre-formed sorting rules and common interpretation. Again, we found more sustained information alignment for the second experiment half. For the first experiment half, late information alignment was present between 510 and 575 ms and between 775 and 825 ms after stimulus onset. For the second experiment half, we found information alignment between 225 and 280, and for much of the time-window between 385 and 925 ms. Together, these results suggest the early information alignment, that was evoked by visual information, did not change over the course of the experiment. However, later alignment, that was unique to real pairs and likely driven by higher order cognitive processes such as attention and/or decision-making, was more persistent over time in later compared to earlier experiment trials.

### 2.4. Beyond shared rules: evidence for pair-specific information alignment

To assess whether this late neural information alignment was driven by representations that were unique to the pair, beyond those driven by shared rules, we conducted an exploratory analysis to compare real pairs to pseudo pairs who shared the same stimulus classification rules. This analysis was conducted separately for the first and second half of the experiment, and was based on 16 of the 24 pairs, as some pairs had no other pairs with the same trial order that used the same rules. Our results show that the late alignment was only partially driven by the dimensions participants used to categorise visual stimuli (Figure 5). Indeed, the late alignment in real pairs was significantly higher than that observed in pseudo pairs where participants used the same dimensions to categorise the stimuli. Stronger alignment was already observed in the first half of the experiment, with a significant cluster from 335 ms to 435 ms, and this effect appeared to strengthen in the second half, with extended significant clusters between 570 ms and 875 ms. Interestingly, this stronger alignment in the second half was observed despite higher correlation also being observed for pseudo pairs, likely reflecting the reinforcement of shared rules. These results suggest that late alignment was driven not only by shared categories collectively formed a priori but also by pair-specific representations emerging from real-time, trial-to-trial interactions through shared perceptions, decisions, collective feedback, and communication between blocks. Furthermore, these results indicate that the strengthening of this late alignment in the second half reflects both non-pair-specific (shared rules) and pair-specific information alignment.

**Figure 5.**
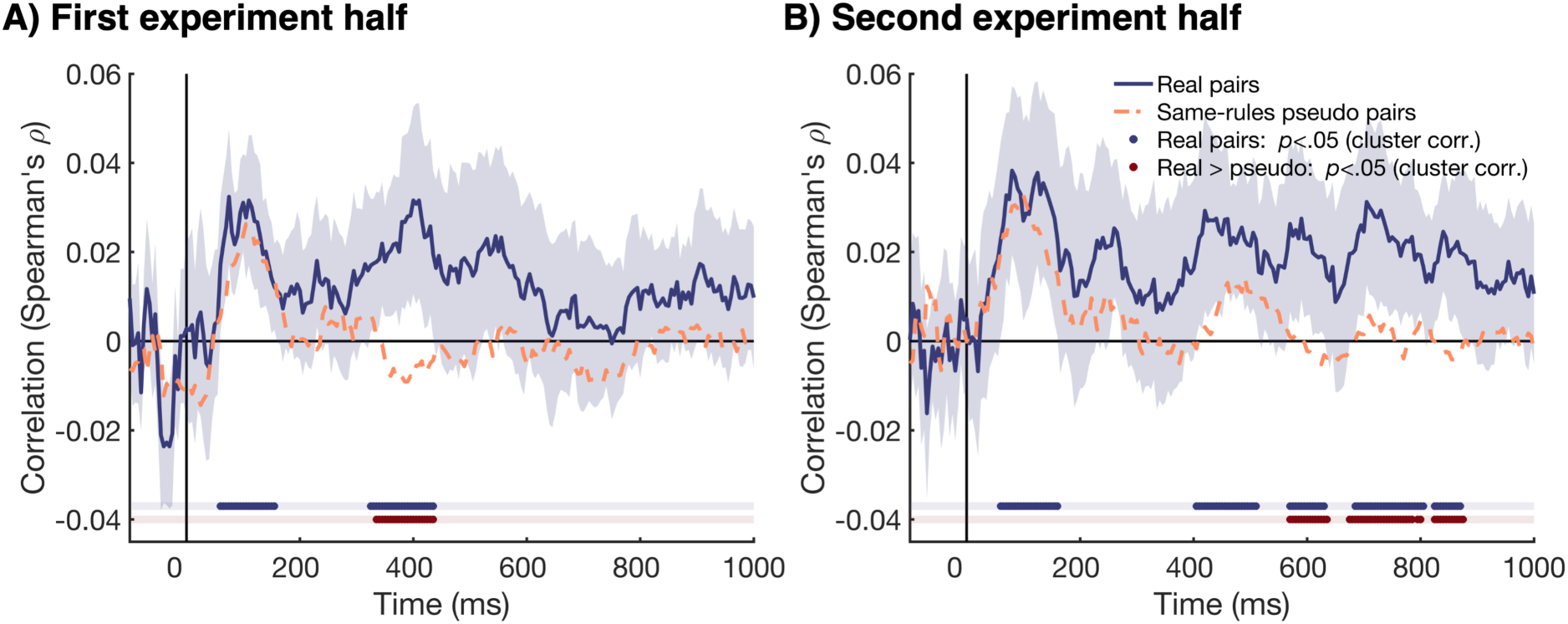
The contribution of shared rules to neural information alignment, for the first (A) and second (B) experiment half. All plotting conventions are the same for figure 5A and 5B. The time-course of information alignment is shown in dark blue. This measure was obtained by collapsing the temporal generalisation matrix of dissimilarity matrix correlations between the two individuals in each pair along one time dimension. The shaded area represents the 95% confidence interval. Significant correlations (cluster-corrected p < .05) are shown in dark blue below. The dashed orange line shows the time-course of correlations for same-rules pseudo pairs, who are randomly matched individuals who independently came up with the same rules. This shows the information alignment that is driven by cognitive processes associated with the same rules, as well as sensory evoked signals. Significant differences between real pairs and same-rules pseudo pairs (cluster-corrected p < .05) are shown in red below, highlighting the socially induced information alignment that cannot be explained by shared rules alone. Note this is based on 16 pairs (32 participants), who were also included in the same-rules pseudo pairs. We excluded 6 pairs because there was no other pair within the same counterbalancing that picked the same categorisation rules.

### 2.5. No evidence for transfer of late information alignment beyond the task

Our results, presented above, show evidence for an alignment in neural information between individuals in a pair during the categorisation task. Importantly, this information alignment was stronger for real pairs compared to random pseudo pairs for the later time-window (∼200 ms onwards). In addition, we found this alignment was, at least in part, driven by pair-specific processes, and it became more sustained as the experiment progressed. To assess whether this alignment was specific to the task, or whether it persisted beyond the categorisation task, we repeated the same analyses on EEG data from a transfer task, which was used as a pre- and post-test. Participants performed the pre- and post-test in the same experimental setup as the main task. The task consisted of a 1-back task on the same stimuli as used in the main categorisation task (Figure 6A). Importantly, the 4 categories that were learned during the categorisation task were not relevant for this task. During this task, participants saw a rapid steam of images: images were presented for 100 ms, followed by a blank screen of 400 ms, followed by the next image. The task of participants was to look for targets in the stream of images. A target was a repeat of the exact same image, twice in a row. The time-course of information alignment during the pre- and post-test is shown in Figure 6A. We found evidence for aligned representations between the participants in the real pairs from 60 to 145 ms after stimulus onset. Importantly, there was no difference between real pairs and pseudo pairs (Figure 6B), suggesting that the alignment was purely driven by visually evoked responses. This means that the neural information alignment found in the categorisation task, that was driven by the integration of shared interpretations, reinforced during the categorisation task, did not persist beyond this task despite participants continuing to share, and perform in, the same environment.

**Figure 6.**
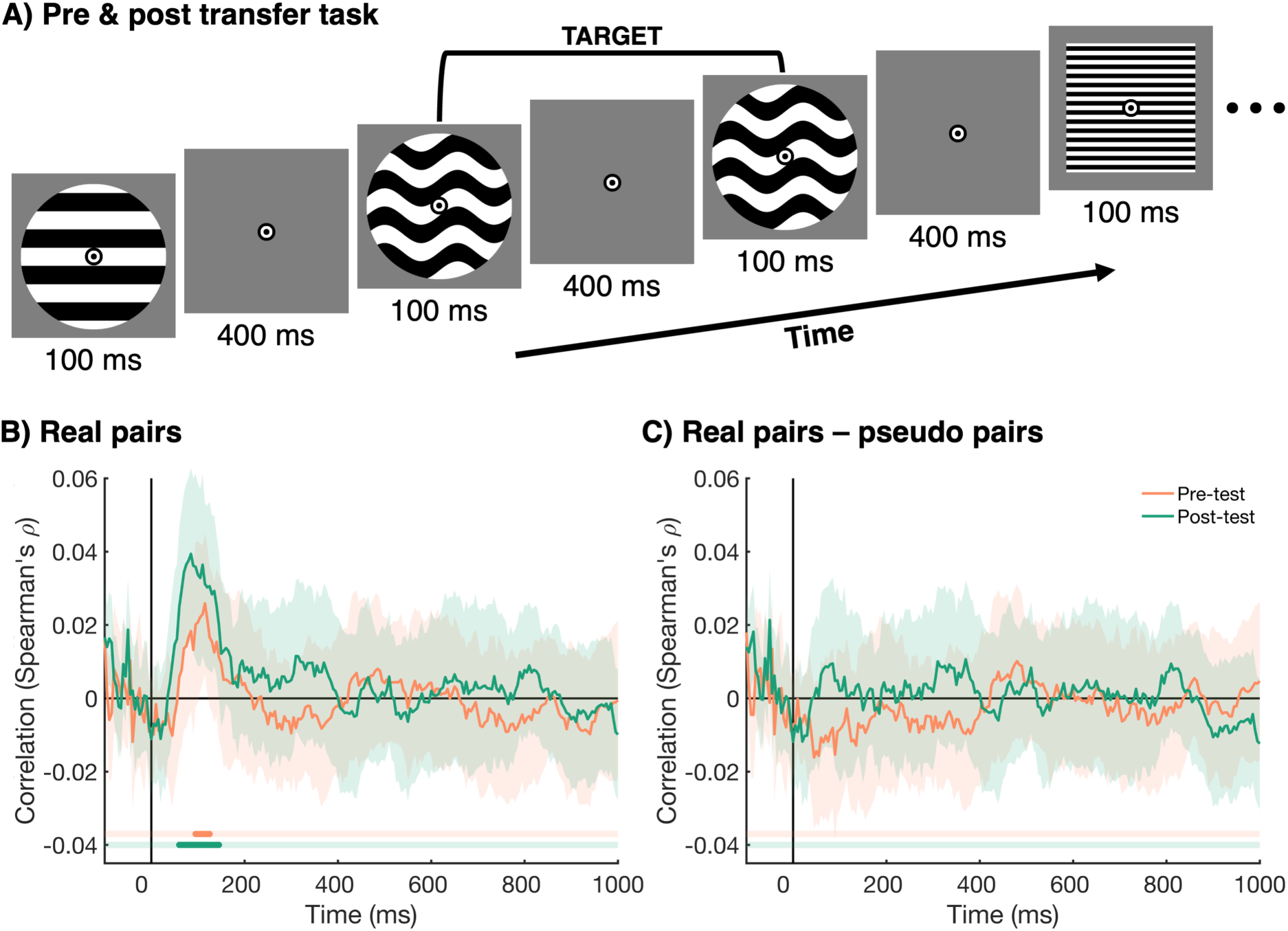
Transfer task (pre- and post-test) design and information alignment. **A)** The same 1-back task was used in a pre- and post-test. In this task, participants saw a stream of images, presented at 2 Hz. The task of the participants was to count the number of 1-back targets (i.e. same image presented twice in a row) for each stream. The agreed upon categories from the main task were not relevant for the pre- and post-test. All plotting conventions are the same as Figure 6B. **B)** The alignment of representations between the two participants in the real pairs for the pre-test (orange) and post-test (green). The shaded areas around the plot lines show the 95% confidence interval, and cluster-corrected *p*-values < .05 are given below the plot in the same colours. **C)** The difference in information alignment between real pairs and pseudo pairs, not constrained by the same rules (real - pseudo pairs). This difference shows the part of the alignment that is not driven by visual information.

## 3. DISCUSSION

In this study, we investigated how social interaction shapes the alignment of neural information between individuals during joint categorisation of visual stimuli. We aimed to distinguish information alignment that was induced by shared rules and interpretations, from visually evoked information alignment due to individuals seeing the same thing at the same time. Pairs of participants performed a categorisation task on stimuli they had first jointly sorted into 4 groups. We used Interbrain RSA on EEG hyperscanning data to get a measure of information alignment between participants in real pairs and pseudo pairs who saw the same stimuli. Results revealed that early information alignment (45-180 ms) was driven by evoked visual responses, as there was no difference between real and pseudo pairs. However, later information alignment (after 200 ms) was stronger for real compared to pseudo pairs. We found that this later information alignment for real pairs increased over the course of the experiment as the shared categorisation rules were reinforced, which was mirrored by an increase in behavioural response similarity. In addition, using pseudo pairs that independently made the same rules, we showed that the later information alignment was not only driven by shared rules, but also by pair-specific representations dynamically strengthened throughout the experiment. Finally, using a separate pre- and post-task, we found that the later information alignment for real pairs did not persist beyond the main task, suggesting it remained largely driven by agreements specific to the task. Together, the results highlight that social interactions play a central role in shaping neural representations in the human brain and that Interbrain RSA is a powerful approach to studying these dynamics using hyperscanning, with promising applications for understanding group collaboration, communication, and decision-making.

### 3.1. Underlying processes of interbrain information alignment

Our results revealed two components behind the interbrain alignment of neural representations emerging during joint categorisation of visual stimuli. First, an early and transient information alignment of large magnitude between 45 and 180 ms driven by shared evoked visual information – seeing the same thing at the same time – as highlighted by the lack of difference between real and pseudo pairs in this timeframe. Importantly, our results also showed that, from 200 ms after stimulus onset onwards, information alignment was stronger for real pairs compared to pseudo pairs. Critically, the pseudo pairs saw the same stimuli in the same order as the real pairs, which means stronger information alignment for real pairs was not driven by shared visual information. Rather, it suggests that this alignment reflects higher-level cognitive processes. An additional analysis, based on pseudo pairs where participants independently picked the same categorisation rules, showed that this late neural information alignment was driven both by non-pair-specific and pair-specific information. These results suggest that higher-level perceptual and decisional processes, related to pre- formed shared categories, drive information alignment that is not pair-specific. Real-time feedback and communication then further refine this alignment during the experiment, producing pair-specific alignment in real pairs.

One possibility is that the non-pair-specific component of the information alignment from 200 ms onwards, at least in part, reflects attentional enhancement of task-relevant features, which is in line with previously reported feature-based attention effects on information coding in the brain, emerging after 200 ms for simple visual stimuli (Moerel et al., 2022, 2024; Smout et al., 2019), novel objects (Goddard et al., 2022), and real-world objects (Grootswagers et al., 2021). An alternative, but not mutually exclusive, explanation is that the later information alignment could represent the 4-way categorisation decision that participants are making together, which is supported by previous work showing that decisions can be decoded from approximately 140 – 180 ms after stimulus onset onwards (Bode et al., 2012; Moerel et al., 2024). Moreover, later alignment was more distributed across the scalp, with stronger contributions of temporal, central and frontal regions, which is in line with brain regions found in previous research to support cooperation during joint activities (Czeszumski et al., 2022; Zhao et al., 2024), and more generally, the encoding of task- and decision-relevant information (Jackson et al., 2016; Jackson & Woolgar, 2018; Moerel et al., 2024; Woolgar et al., 2015). Further evidence that late alignment was driven by task-related cognitive processes, such as attention and/or decision-making, comes from the transfer task. In this task, participants saw the same stimuli as in the main categorisation task, but critically, participants did not have to group the stimuli into the 4 categories, which means the previously agreed upon rules were no longer relevant. Here, we found no difference in information alignment between the real and pseudo pairs, suggesting that this alignment was only driven by shared visual input. This finding adds further evidence that the late alignment, observed for the categorisation task only, was indeed driven by processes beyond shared visual input, and likely mediated through the agreed upon rules. In contrast to early visually evoked alignment, the late alignment observed during the categorisation task was found to be a dynamic process that was reinforced over time, as evidenced by stronger magnitude in the second half of the experiment compared to the first one. Importantly, this finding fits with the behavioural data, which indicated stronger behavioural similarity, as well as increased accuracy, as the experiment progressed. Together, these results suggest that late alignment was not static from the start of the experiment but increased over time as the shared rules were reinforced with repeated practice.

Finally, there was a pair-specific component to the late alignment, with stronger alignment for real pairs compared to pseudo pairs made up of people who independently chose the same sorting rules. This suggests that the late alignment is only partly driven by information related to the sorting rules and processes, such as attention to the relevant stimulus features and decision-making related to the categories. This alignment also likely reflects further refinement of higher-level cognitive processes that uniquely emerge within real pairs through real-time interactions. On each trial, participants received feedback at the level of the pair. This means that participants had information not only about their own accuracy, but that of their partner as well. In addition, pairs were in the same room and were allowed to talk during the breaks, with a total of 19 breaks spread throughout the task. These breaks allowed participants to discuss and potentially refine their strategies, informed by the shared feedback they received. This could have resulted in strategies that were unique to the pair. This explanation fits with our finding of more sustained alignment, driven by pair-specific information, during the second compared to the first half of the experiment. This might have also contributed to behavioural alignment increasing over the course of the experiment. Together, these findings suggest that the observed late alignment reflects not just processes related to shared rules, but also the emergence of shared, pair-specific processes shaped by ongoing interaction and joint feedback throughout the task.

Our findings reveal that when rules are agreed upon and actively upheld, it reinforces the way information is represented and aligned across individuals’ brains. This process may have significant implications for group behaviour and decision-making, potentially contributing to the formation of lasting traditions and social norms (Cialdini & Goldstein, 2004; Jacoby et al., 2024; Tomasello et al., 2005). Future research could explore how different types of social agreements, beyond categorisation rules, enhance information alignment between individuals. Additionally, studies using longer time scales, or possibly even cross-generational approaches, could assess the durability of social information alignment over time, and whether this is the same across different cultures.

### 3.2. Interbrain RSA: a novel method for hyperscanning research

Our results highlight the exciting potential of Interbrain RSA for studying neural representations and interbrain alignment during joint activities (Coey et al., 2012; Keller et al., 2014; Marsh et al., 2009; Sebanz et al., 2006). By moving away from brain activation, and instead indexing information content, Interbrain RSA effectively captures what information is driving the alignment between brains, and it therefore has the potential to distinguish between subtle sensory and cognitive processes and their convergence across individuals. By focusing on the converging information contained across neural signals, Interbrain RSA provides new insights into neural representations and their underlying mechanisms supporting everyday social behaviours. Varlet and Grootswagers (2024) previously demonstrated the power of Interbrain RSA in capturing information alignment driven by shared visual inputs through EEG hyperscanning simulations. In this study, we show for the first time the method’s effectiveness in capturing not only sensory but also social information alignment in real-time social interactions. Using Interbrain RSA, we were able to capture the information alignment that uniquely emerged from the social interaction itself, over and above the alignment from shared visual input, which in turn increased similarity in behavioural responses in participant pairs. This new method has the advantage of directly capturing the neural information alignment of both lower- and higher-level representations, including their underlying spatiotemporal dynamics, providing a deeper understanding of the sensory and cognitive processes converging between individuals during social interaction (Czeszumski et al., 2020; Dumas et al., 2010; Holroyd, 2022; Varlet et al., 2020). An additional advantage of Interbrain RSA, over traditional interbrain synchrony measures, is that it provides a single, experiment-level estimate of information alignment between brains. Traditional synchrony analyses are highly sensitive to moment-to-moment fluctuations, such as shared environmental noise. However, Interbrain RSA aggregates information across the entire experiment, making it robust against transient artefacts. The classifier is trained and tested at different times in the experiment, and only signals that reliably inform the classes of interest are used by the classifier. This allows researchers to isolate genuine information alignment between participants, rather than effects driven by being in the same environment at the same time. Whereas the Interbrain RSA method provides a powerful tool to investigate information alignment between brains, its validity ultimately depends on careful design. It is essential to control for confounding factors that might co-vary with the conditions of interest and to ensure that structure of the representational matrix is sufficiently sensitive to the effect being investigated.

In future research, Interbrain RSA could provide valuable insights into group collaboration, communication, and how shared cognitive processes emerge in various social contexts (D’Ausilio et al., 2015; MacRitchie et al., 2017; Marsh et al., 2009; Miles et al., 2023; Sebanz et al., 2006; Speer et al., 2024). While in this experiment the late information alignment was driven by the rules the pairs agreed upon, unique ways of interpreting or representing the environment can also emerge within a group through the group’s specific preferences, decisions, or traditions (Cialdini & Goldstein, 2004; Jacoby et al., 2024; Konvalinka et al., 2011; Néda et al., 2000; Tomasello et al., 2005). These shared ways of interpreting the world could drive performance in teams and communities by enabling them to act more effectively together, as they interpret and respond to events into their environment under a common set of rules. While Interbrain RSA has not been used yet in other perceptual modalities, classic RSA has been successfully used to investigate the perception of sound (Su et al., 2014), touch (Lee Masson et al., 2018), and even smell (Fournel et al., 2016), highlighting the potential of this method to assess the alignment of perceptual information in and across different modalities. Notably, by abstracting from brain activity into meaningful representational structures, Interbrain RSA can effectively index and compare neural representations across different brain regions and different time scales when using interbrain representational connectivity and temporal generalisation analyses as shown here. In addition, it provides the freedom to compare neural representations across neuroimaging modalities and tasks. These methodological advances offer promising avenues for deepening our understanding of the neural processes underlying real-time social interactions, which are often disrupted in mental illnesses like schizophrenia, autism, and social phobia phobia (Charman, 2003; Lord et al., 2020; Moreau et al., 2024; Raffard et al., 2015; Varlet et al., 2012, 2014).They could lead to new insights into key markers of these conditions.

### 3.3. Conclusion

In this study, we used Interbrain RSA, a novel method, to reveal for the first time that information content aligns between individuals’ brains as they collaborate. Importantly, this alignment goes beyond basic visual processing from shared stimuli to include late information alignment, which begins around 200 ms after stimulus onset. This information alignment is driven by shared rules and associated higher-level cognitive processes, such as enhanced collective attention to the two features needed to group the stimuli and decisions about the which group a stimulus belongs to. It also reflects pair-specific information beyond shared rules, likely emerging from real-time interactions through shared feedback and communication. This late alignment between participants strengthened over time, alongside an increase in behavioural similarity as the task progressed, indicating that shared cognitive frameworks develop through deeper engagement in collaboration. Critically, this late alignment did not persist in a subsequent task where the agreed upon rules no longer applied, highlighting that such interbrain alignment depends on the active maintenance of relevant rules and common interpretations. Interbrain RSA opens new pathways for understanding how individuals collaborate. Establishing shared rules may promote a unified interpretation of the environment, enhancing team performance by enabling coordinated interpretation and response to environmental demands.

## 4. MATERIALS & METHODS

### 4.1. Participants

Fifty-two participants took part in the EEG experiment (26 pairs). Two pairs of participants were excluded from the analysis due to an issue with the counterbalancing of the trial order, resulting in a final sample of 24 pairs (27 female/25 male, 48 right-handed/4 left-handed, mean age = 23.58, SD = 2.78, age range = 19 - 31). The sample size was based on publicly available data from previous work that introduced the Interbrain RSA method and reported the difference in information alignment between same vs. different object viewing across 10,000 simulations (Varlet & Grootswagers, 2024). We calculated the effect size for each simulation and used the median effect size to determine the power for this study. We estimated that a 1-tailed t-test with 12 pairs would yield 80% power, and 19 pairs would yield 95% power. We tested 24 pairs to ensure sufficient power even for smaller effects. Pairs consisted of both same-sex and different-sex pairs. All participants reported normal or corrected-to-normal vision. The study was approved by the ethics committee of Western Sydney University, and participants provided both written and verbal consent before participating. The EEG session took approximately 2 hours to complete, and participants received AU$30/hour for their participation.

### 4.2. Stimuli and experiment procedure

The two participants in the pair were seated back-to-back, each facing a computer screen showing the same visual input. The EEG experiment consisted of a main task, in which participants performed a stimulus categorisation task. There were also a pre- and post-test that served as a transfer task. The same stimuli were used across all experimental parts and consisted of 16 greyscale images varying on 4 dimensions (Figure 1B), with 2 levels per dimension: spatial frequency (thick vs. thin lines), pattern (wavy vs. straight lines), contrast (high vs. low) and general shape (circle vs. square). All stimuli were presented at 200 by 200 pixels (approximately 5.06 by 5.06 degrees of visual angle) on a mid-grey background on a VIEWPixx/EEG monitor (VPixx Technologies Inc., Saint-Bruno, Ontario, Canada) running at a refresh rate of 120 Hz. The experiment was programmed using the Psychopy library (Peirce et al., 2019) in Python. The stimuli were always presented behind a central fixation bullseye (18 by 18 pixels, approximately 0.46 by 0.46 degrees of visual angle) (Figure 1C and Figure 5A), and participants were instructed to maintain fixation on the bullseye throughout the experiment.

#### 4.2.1. Categorisation task

The stimulus categorisation task formed the main part of the EEG experiment (Figure 1C). First, participants saw all 16 images presented on the screen. They were asked to work together to sort the images into 4 groups of 4 stimuli, by dragging and dropping them to the parts of the screen labelled ‘Group 1’ (top left), ‘Group 2’ (bottom left), ‘Group 3’ (top right), and ‘Group 4’ (bottom right) (Figure 1C, top panel). To do this task, participants had to select two of the stimulus dimensions while ignoring the other two dimensions. For example, if the pair selected spatial frequency and shape, while ignoring pattern and contrast, the 4 groups would be thick circles, thick squares, thin circles, and thin squares. Participants were allowed to talk to each other during this part. They could both move the stimuli and were able to observe the movements of the other person as their computer screens showed identical displays. Both participants had to confirm they approved the selected groups before the second part of the categorisation task commenced. The EEG data recorded during this first part were not analysed.

After the participants in the pair jointly made the 4 groups, participants engaged in a joint categorisation of the visual stimuli. As we analysed the EEG data from this part, participants were asked not to talk during this part, but they were allowed to talk during the breaks. There were a total of 19 breaks, spaced every 32 trials, with less than 3 minutes between breaks. In this task, participants saw a stimulus for 100 ms, followed by a blank screen for 900 ms. The task of the participants was to categorise the image based on the 4 groups the pair previously made together (Figure 1C, bottom panel). Both participants in the pair were asked to respond. Participants used 4 buttons on a QWERTY keyboard to respond, marked with white dots. One of the participants used the ‘Q’, ‘W’, ‘O’, and ‘P’ keys, and the other participant used the ‘A’, ‘S’, ’K’, and ‘L’ keys. The mapping of the response buttons changed on each trial, and was shown on the screen for 1000 ms, during which the participants could respond. The order of the numbers on the response screen corresponded to the order of the buttons. For example, if the image was from ‘Group 1’, and the response mapping was ‘3 – 2 – 4 – 1’, this means the rightmost button was correct. The response button mapping screen served two purposes: 1) it introduced a delay between the stimulus and response, which allows us to analyse the part of the EEG signal (0 – 1000 ms) that is not contaminated by motor execution, and 2) it made the response-button mapping unpredictable, preventing participants from preparing a motor response during the time window used in the analysis. If participants did not respond within 1000 ms, the experiment would automatically continue. After the response time-out was reached, participants received feedback on whether they were correct, which was presented on the screen for 500 ms. We determined the response accuracy for each participant in the pair by verifying that the responded group number matched their previously chosen group. The fixation bullseye turned green if both participants were correct, orange if one of the participants was correct and the other incorrect, and red if both participants were incorrect. The same feedback was always presented to both individuals in the pair, which means that in the case of one correct response, it was not specific about which individual made the error.

The categorisation task consisted of 20 blocks. At the start of each block, participants saw a reminder of the 4 groups of 4 they had made previously. A single block contained 32 trials, with 2 repeats of the 16 unique images. The images were presented in a random order, and all 16 images were presented before any repeated images occurred. Participants were offered breaks in between each block, and both participants had to confirm before the next block was started. This task took approximately 30 to 45 minutes to complete, depending on breaks.

#### 4.2.2. Transfer task (pre- and post-test)

Pre- and post-test in this task were used to determine whether interbrain information alignment persisted beyond the main categorisation task. In this task (Figure 5A), participants saw a rapid stream of images, presented at 2 Hz, and looked for a presentation of the exact same image twice in a row. Each image was presented for 100 ms, followed by a blank screen of 400 ms, followed by the next image. To avoid a manual response during the stream of images, participants were asked to count the number of targets (1-back repeats) in the stream. At the end of each stream, they were asked to report the number of targets.

There were 1 – 3 possible targets in a stream. For each target event, the two target (i.e., same) stimuli and two additional consecutive stimuli, that were added as padding, were removed from analysis. A single stream consisted of 164 to 172 image presentations, depending on the number of targets in the stream. This leaves 160 stimulus presentations that were included in the analysis for each stream, consisting of 10 repetitions of the 16 unique images. There were three streams of images, and participants were offered breaks between each stream. The pre- and post-test took approximately 5 minutes each to complete

#### 4.2.3. Counterbalancing

To be able to distinguish visually evoked neural information alignment from alignment that was not driven by shared visual information, we compared Interbrain RSA between real pairs and randomly matched-up pseudo pairs who did not perform the task together but saw the same stimuli. To ensure that any difference between the real pairs and randomly matched pseudo pairs was not driven by the stimulus order, we used the same counterbalancing of the trial order for multiple pairs. There were 4 unique trial orders, each seen by 6 pairs of participants. Any reference to randomly matched pseudo pairs refers to participants who saw the same trial order.

### 4.3. EEG acquisition and pre-processing

We recorded continuous EEG data from 64 channels, digitised at a sampling rate of 2048 Hz, using the BioSemi active electrode system (BioSemi, Amsterdam, The Netherlands). We used the international standard 10-20 system to place the electrodes (Oostenveld & Praamstra, 2001). We used the Fieldtrip toolbox in MATLAB to pre-process the data (Oostenveld et al., 2011). We re-referenced the data to the average of all channels, applied a high-pass filter of 0.1 Hz, low-pass filter of 100 Hz, and band-stop filter for 50 Hz line noise. We down-sampled the data to 200 Hz and created epochs from -100 ms to 1000 ms relative to stimulus onset. We applied a baseline correction using the -100 to 0 ms as the baseline. No further pre-processing was applied. Note the selected epoch only included the part of the trial before the button mapping was presented and therefore excluded the part of the trial where motor preparation and execution could occur.

### 4.4. Behavioural analysis

We calculated two behavioural measures for the categorisation data. First, for each block in the experiment, we calculated the accuracy, separately for both individuals in the pair. Second, we calculated the proportion the same response was given by both individuals, regardless of whether the response was correct or incorrect. This resulted in a single behavioural alignment score per pair for each block. For both the behavioural accuracy and behavioural alignment measure, we excluded trials on which one or both participants did not provide a response before the time-out was reached (1000 ms).

### 4.5. Interbrain Representational Similarity Analysis

We used Interbrain RSA (Kriegeskorte et al., 2008; Varlet & Grootswagers, 2024) to assess the degree to which representations were aligned between the two individuals in each pair during the categorisation task. First, for each participant separately, we applied temporal smoothing to the data with a kernel of 20 samples (100 ms). Then, we created a Representational Dissimilarity Matrix for each time-point in the epoch. The dissimilarity matrix reflects the dissimilarity in the pattern of activation across all 64 channels for each combination of two stimuli. We used decoding accuracy as a measure of dissimilarity, with higher decoding accuracy reflecting a more dissimilar pattern of activation driven by the two stimuli. The 16 unique stimuli were repeated 40 times during the categorisation task, and all 16 images were presented before any repeated images occurred. We used a Linear Discriminant Analysis classifier, with a 40-fold cross-validation. We did this for each combination of the 16 unique stimuli, resulting in a 16 by 16 dissimilarity matrix for each time-point and participant.

To determine the information alignment between the two individuals in the pair, we correlated the dissimilarity matrices of the two individuals. To capture information alignment that emerged at different times between participants in the pair, we used a temporal-generalisation approach, where we calculated the Spearman correlation between the dissimilarity matrices of the two individuals for each combination of time-points in the epoch. We averaged the time-by-time matrix of each pair with a permuted version, where the x and y axes are flipped, to make it symmetrical around the diagonal. In addition to performing this analysis for the real pairs of participants, who did the task together, we repeated the same analysis for the randomly matched pseudo pairs. Importantly, the matched trial orders ensured that the individuals in the pseudo pairs saw the same stimuli in the same order. This allowed us to determine whether the shared representations were driven by visual stimulus information. Because the individuals in the pseudo pairs did not perform the grouping task together, the agreed upon rules were likely different between individuals. For each of the 10,000 iterations, we made 24 random pseudo pairs and calculated the correlation between the two individuals in each pseudo pair, for each combination of time-points. We then averaged across the 10,000 iterations. Note that the order of the time-points was kept intact, and no shuffling across time-points was performed. To condense the time-by-time matrix of interbrain information alignment into a time-course, we averaged over one of the time dimensions of the symmetric time-by-time matrix. We did this for both the real pairs and pseudo pairs. This resulted in a single time-course, that captures shared representations that occur at different points in time for the two participants in the pair. We repeated the same analysis described here separately for the first and second half of the experiment, to assess whether the shared representations become more persistent over time, when individuals perform the task together.

To determine whether observed alignment was driven by shared rules alone, or whether there was pair-specific alignment beyond shared rules, we ran an exploratory analysis in which we formed pseudo pairs based on the categorisation rules that each pair picked. We excluded 6 pairs for this analysis, as there was no other pair within the same counterbalancing that picked the same categorisation rules, resulting in a sample of 16 pairs (32 individuals) for this analysis. The individuals could be matched-up into 44 same-rules pseudo pairs, with each individual contributing to multiple pseudo pairs. For each same-rules pseudo pair, we calculated the alignment between individuals as described above, separately for the two experiment halves. We correlated the dissimilarity matrices for each combination of time-points in the epoch and then collapsed along one time dimension. To be able to compare the real and same-rules pseudo pairs, we also re-calculated the alignment for the real pairs, by only including the selected 16 pairs from the total sample of 24 pairs. Higher correlation in real compared to same-rules pseudo pairs enables capturing alignment driven by information beyond shared categorisation rules.

Finally, we determined whether any information alignment, found during the categorisation task, persisted beyond the main task. To this end, we repeated the analysis described above using the data from the transfer task (pre- and post-test).

### 4.6. Spatial dynamics of interbrain alignment

We used a searchlight at the channel level to determine which channels had the strongest contribution to the information alignment for the categorisation task. For each of the 64 channels, we selected 4 or 5 neighbouring channels, and repeated the analysis described above. This resulted in an alignment measure for each combination of two channels, for all combinations of time-points. We averaged across one of the time dimensions in the same way as described above. In addition, we averaged across one of the channel dimensions. This resulted in a topography, that reflected for each channel and each time-point how strong the alignment is with any time-point and channel of the other individual in the pair. For plotting purposes, we averaged the topographies within 100 ms time bins. We did the same for the randomly matched pseudo pairs (10,000 iterations).

The searchlight measure provides insight into the spatial pattern of the information alignment. However, it cannot inform us about the combination of regions between brains that drive the alignment. Therefore, we calculated an additional spatial measure. We examined which clusters of channels contributed most strongly to the shared neural representations between participants by repeating the whole-brain analyses described above for individual clusters of channels. We excluded all channels on the midline, and divided the remaining channels into 6 clusters: Frontal Left (Fp1, AF3, AF7, F1, F3, F5, F7), Frontal Right (Fp2, AF4, AF8, F2, F4, F6, F8), Temporal/Central Left (FC1, FC3, FC5, FT7, C1, C3, C5, T7, CP1, CP3, CP5, TP7), Temporal/Central Right (FC2, FC4, FC6, FT8, C2, C4, C6, T8, CP2, CP4, CP6, TP8), Parietal/Occipital Left (P1, P3, P5, P7, P9, PO3, PO7, O1), and Parietal/Occipital Right (P2, P4, P6, P8, P10, PO4, PO8, O2). We obtained a single dissimilarity matrix for each of the 6 channel clusters, for each time-point and participant. We determined two time bins of interest: early (45 – 150 ms after stimulus onset) and late (200 – 1000 ms after stimulus onset). We then correlated the dissimilarity matrices for the two participants in the pair for each combination of channel clusters, for all the time-points within each window of interest. We did the same for the randomly matched pseudo pairs (10,000 iterations). By collapsing across regions and time-windows of interest, we were able to assess information alignment between all the combinations of channel clusters.

### 4.7. Statistical inference

We used *t*-tests to statistically assess 1) whether information alignment within a pair was greater than chance, and 2) whether information alignment within a pair was greater than that for randomly matched pseudo pairs. We ran a single 1-tailed *t*-test for each time-point in the time-course of information alignment, obtaining a single *t*-value for each time-point. To correct for multiple comparisons, we used cluster-corrections (Maris & Oostenveld, 2007; Stelzer et al., 2013). We randomly sign-flipped the *t*-values and obtained the clusters where all consecutive *t*-values were above the critical value of 1.711. The order of the time-points was kept intact to preserve the temporal structure of the data. For each observed cluster, we calculated the sum of all *t*-values in that cluster. We then selected the largest observed sum of *t*-values. We repeated this 10,000 times, obtaining 10,000 measures for the largest cluster observed when *t*-values were randomly sign-flipped. We then selected the 95^th^ percentile as a threshold. Only real clusters with a summed *t*-value above the threshold were selected. We repeated this analysis for 1) the time-course of correlations for the full categorisation experiment, 2) the time-course of correlations for the first and second half of the categorisation experiment separately, and 3) the time-course of correlations for the pre- and post-test.

## ACKNOWLEDGMENTS

We thank Louise Daix-Moreux for her help with data collection. This work was supported by an Australian Research Council (ARC) Discovery Project awarded to M.V. (DP220103047) and an ARC Discovery Early Career Researcher Award awarded to T.G. (DE230100380).

## **5.** DATA AND CODE AVAILABILITY

The analysis code, figures, and an example dataset can be found via the Open Science Framework (https://doi.org/10.17605/OSF.IO/HE4TU). The full dataset will be made available upon publication.

